# Reversal of carbapenem resistance in *Pseudomonas aeruginosa* by camelid single domain antibody fragment (VHH) against the C4 dicarboxylate transporter

**DOI:** 10.1101/2023.06.20.545680

**Authors:** Anil Kumar Nagraj, Manjiri Shukla, Mansi Kulkarni, Pratik Patil, Mrunal Borgave, Sanjiban K Banerjee

**Affiliations:** AbGenics Life Sciences Pvt. Ltd. Pune, India 411045

**Keywords:** VHH camelid antibody, *P*. *aeruginosa*, carbapenem resistance, meropenem, phage display, combination therapy

## Abstract

The COVID-19 pandemic has hastened the problem of nosocomial drug-resistant pathogens and is exerting a huge toll on hospitalized patients in critical care settings. Most small molecule antibiotics are susceptible to bacterial resistance mechanisms and constantly becoming ineffective leading to rapid shrinkage of the antibiotic armamentarium. *Pseudomonas aeruginosa*, one of the most common pathogens in hospital-born infection and considered critically dangerous by WHO and CDC, is extremely difficult to treat with frontline drugs, the carbapenems. In this study, we developed a camelid antibody fragment (VHH) library against whole *P. aeruginosa* and isolated a highly potent neutralizing hit (PsC23) that selectively targets *P. aeruginosa .* At 25 µg/mL, PsC23 inhibited growth of the ATCC 27853 and a locally isolated carbapenem resistant MCC 50428 *P. aeruginosa*. The target of PsC23 is the C4 dicarboxylate transporter that transports C4 metabolites to the glyoxylate shunt during oxidative stress that is present in pathogens but not the human host. This ultimately results in the blockade of the shunt affecting bacterial energy transduction that leads to disruption of drug efflux. Interestingly, in a neutropenic mouse with MCC 50428 systemic infection, PsC23 in combination with meropenem completely reversed the drug resistance and eliminated the pathogens from the blood. PsC23 was stable in human serum and had no hemolytic or cytotoxic effect on human cells. Taken together, this VHH if co administered with the currently available carbapenems would reverse carbapenem resistance and could be used to effectively control *P. aeruginosa* in critical care settings.

## Introduction

Worldwide, infection due to *Enterococcus faecium, Staphylococcus aureus, Klebsiella pneumoniae, Acinetobacter baumannii, Pseudomonas aeruginosa* and *Enterobacter spp* (ESKAPE) group that are multidrug-resistant (MDR) pathogens in critical care settings have emerged as a new health care crisis (1). In 2013, the Centers for Disease Control and Prevention and World Health Organization (2017) listed these pathogens as a serious threat. Among them, the carbapenem resistant *Enterobacteriaceae, Pseudomonas,* and *Acinetobacter* were placed in the highest threat priority category. Some of the *P. aeruginosa* strains are resistant to nearly all antibiotics, including carbapenems (2) and are common in hospitalized patients especially the immuno-compromised with a lung and urinary tract infections, skin infections in burn patients, diabetic ulcer and cystic fibrosis that cause increased morbidity and mortality due to suboptimal efficacy and cross-resistance to multiple drugs (3,4,5,6). The occurrence of the antibiotic-resistant *Pseudomonas* is more predominant in hospital settings in India and developing countries as compared to the industrialized nations of Europe and North America (7,8,9).

Carbapenems (meropenem and imipenem) are frontline drugs that have been widely used against many drug-resistant bacterial infections (10). Although carbapenems are widely used for treating *P. aeruginosa*, the prevalence and surge of resistant strains in recent years have made them less effective which necessitates the need to develop newer therapeutic options (11) as a result of which novel approaches including antimicrobial peptides, phage therapy, vaccines and antibodies have been utilized against MDR-pathogens (12).

Camelids (camels, llamas, and alpacas) produce unconventional single heavy chain antibodies that are devoid of light chain (13). Despite their small size (one-tenth) than conventional antibodies, they have a high affinity for antigens due to their unique structural attributes (14). They have additional advantages over conventional antibodies such as higher thermal and chemical stability and ease of production by simple microbial systems and have enabled their use for therapeutic and diagnostic applications (15). VHH fragments have been exploited as drug candidates against many pathogens and parasites, such as *Listeria monocytogenes* (16), *Leishmania infantum* (17) and Coronavirus (18). Moreover, Caplacizumab, the camelid nanobody against human anti-von-Willebrand factor is already in the market as an approved drug having obtained regulatory approvals in Europe and USA highlighting the potential of VHHs as therapeutic molecules (19). In this study, we constructed a VHH fragment library against *P. aeruginosa* from an immunized Indian dromedary camel using the phage display and isolated a neutralizing antibody fragment VHH hit (PsC23). The potential target of this hit was identified by mass spectrometry was revealed to be a component of the C4 dicarboxylate transporter. We characterized the target by microbiological and biochemical experiments and evaluated the anti-bacterial activity of PsC23 against a clinical carbapenem resistant *P. aeruginosa* MCC 50428 *in vitro* and studied its therapeutic efficacy in combination with carbapenems in a neutropenic mouse model with systemic infection where a complete clearing of the infection was observed validating PsC23 as a lead candidate for further development as a therapeutic molecule.

## Materials and methods

### Immunization of camels and construction of the *P aeruginosa* VHH library in *E. coli*

A 7-10 year old Indian dromedary camel (*Camelus dromedarius*) was infected with an inactivated meropenem-resistant *P. aeruginosa* (strain MCC 50428, National Centre for Microbial Resources, Pune), as described previously (20) with suitable modifications. Briefly, the overnight grown culture was kept for inactivation in PBS containing 0.4% formaldehyde at 4°C overnight. A mixture (1:1 *v/v*) of inactivated *P. aeruginosa* (∼1 x 10^9^ CFU) and alum mineral oil adjuvant (MONTANIDE^TM^ ISA 206 VG, Seppic Inc. USA) was injected subcutaneously on 1^st^, 30^th^ and 45^th^ day. Heparinized blood (60 mL) was collected from the jugular vein on 60^th^ day. Peripheral blood mononuclear cells (PBMCs) were isolated by density-gradient centrifugation (5000 rpm) using Ficoll-Histopaque 1077 (Sigma-Aldrich, USA) and a buffy coat was collected from the interface (21). Total RNA was extracted from PBMCs using Hi-Pure A miniprep RNA extraction kit (HiMedia, India) and converted to cDNA (1 µg) with oligo-dT primers followed by second-strand synthesis using the camelid antibody-specific primers. The amplified VHH fragments were then digested with *SfiI* (NEB, UK) and ligated between *Bgl1*/*Sfi1* site into the pADL-23c phagemid (Antibody Design Lab, USA) and transformed in *E. coli* ER2738 (NEB, UK) by electroporation (2000V 5 ms pulse). The library size was determined by plating various dilutions on 2X YT agar containing carbenicillin (100 µg/mL) and 2% glucose. 10 randomly selected colonies were subjected to colony PCR to check the presence of the VHH insert (Fig. 1a). The library was then stored at −80°C in LB + 50% glycerol till further use.

**Fig. 1.**
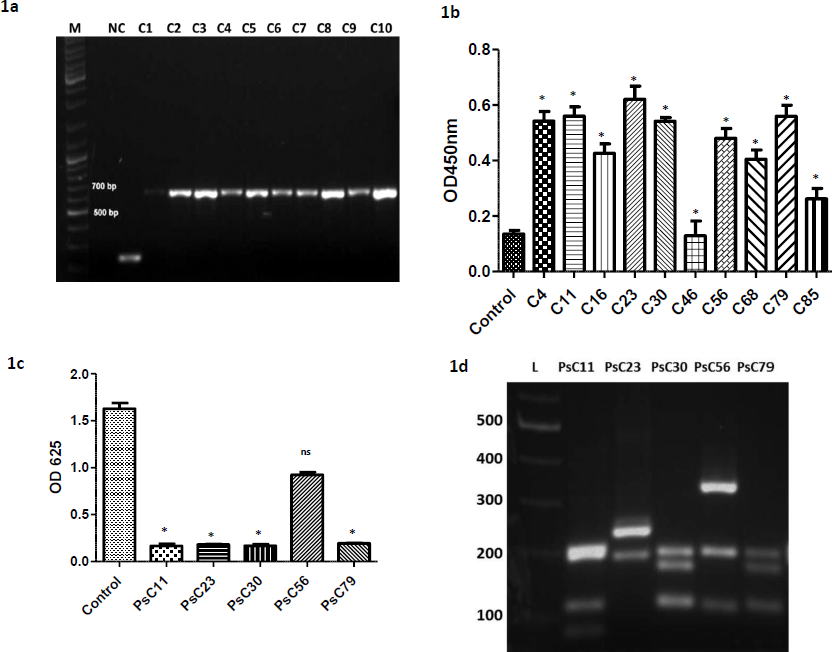
**A.** Agarose gel electrophoresis profile of the VHH fragments amplified by PCR from randomly chosen *E coli* clones from CIL-Ps library cloned in pADL-23c vector. M: Molecular weight marker, NC: (Negative control) is the empty vector, and C1-C10: clones containing the intact VHH fragments. **B.** Whole cell phage-ELISA with supernatant of induced *E coli* clones containing the full-length VHH in pADL-23c vector. Clones were categorized by the fold increase in binding compared to control by compared to negative. A more than 4-fold increase was considered as a strong binder. Low-binding hits (< 4 fold binding) were not progressed. C23 showed significantly greater OD than control (\**P*<0.0001) and was referred to as PsC23 subsequently**. C.** Neutralization of *P. aeruginosa* with the supernatant of induced *E. coli* secreting anti-Pseudomonas VHH fragments from pADL vector as quantitated by measurement of OD 625 after 16 hours. PsC23 significantly inhibited the growth of *P.aeruginosa* compared to control (\**P*<0.0001). **D.** RFLP analysis of PCR extracted VHH genes from the hit clones, separated after *BstN1* digestion on 1.5% agarose gel.

### Enrichment and selection of *P. aeruginosa* binding VHH hits by phage display

Enrichment of phages expressing hits was carried in immunotubes (Thermo Scientific, USA) pre-coated with whole-cell *P. aeruginosa.* Three consecutive rounds of bio-panning enriched the population of Pseudomonas-specific binding phages. These phages were allowed to infect *E. coli* ER 2738 and screened for antigen binding hits by phage-ELISA using anti-M13-HRP conjugated antibodies (22). Briefly, *E. coli* ER 2738 colonies were picked randomly and inoculated in 2X YT medium containing carbenicillin (100 µg/mL) in a 96-deep well plate and incubated at 37^0^C, 500 rpm. At 0.4 OD, helper phage M12K07 (5 x 10^11^ pfu/mL) was added with kanamycin (50 µg/mL) and incubated overnight at 30 °C, centrifuged at 500 rpm and phage particles in the supernatant were collected by centrifugation the following day. Meanwhile, whole *P. aeruginosa* MCC 50428 (1 x 10^6^ CFU per well) were coated in ELISA plates (NEST, China) using bicarbonate buffer, pH 9.2, and blocked with 5% skimmed milk (1 h, 37 °C). The wells were washed thrice by PBS with 0.01% Tween-20 (PBST, pH 7.2). 100 µL of the phage supernatant was then added to each well followed by incubation for 1 h at 37°C. Wells were washed with PBST thrice, anti-M13-HRP conjugated antibody (0.4 µg/mL, Sino Biological, USA) was then added followed by 1-hour incubation at room temperature. After washing with PBST, 0.5 mM TMB (Sigma-Aldrich, USA) was added and incubated at RT for 5-15 min till desired blue color was developed. The reaction was terminated by 1% sulfuric acid and optical density (OD) of 450 nm was measured.

### Identification of *P. aeruginosa* neutralizing VHH hits by leaky expression assay

Binding hits having a high reading in phage-ELISA compared to negative control were further evaluated for their neutralization ability against *P. aeruginosa*. Briefly, these colonies were grown in 96-deep well plate under vigorous shaking in the medium supplemented with 0.5% Triton X100 and 0.5% of Glycine to stimulate the release of soluble VHH fragments from the periplasmic space into medium supernatant. The supernatant was passed through a 0.2 µm filter to remove the *E. coli* from the supernatant, diluted twice with LB broth containing 1 x 10^3^ CFU/mL of *P. aeruginosa,* and incubated at 37°C for 24 h and growth was monitored. The clones that distinctly inhibited the growth of *P. aeruginosa* were monitored visually and few leads were identified by OD measurement and picked up for further analysis. The distinctiveness of these clones was analyzed by RFLP by digesting them with *BstN1* (NEB, UK). One of the clones (PsC23) which showed consistent neutralization and strong binding in ELISA was sub-cloned into *E. coli* expression vector pET-28c (+) (Novagen, USA).

### Expression and purification of PsC23 VHH fragment

Overnight grown culture of E. *coli* C41(D3) containing pET-28c with PsC23 was sub cultured in a fresh media and grown till 0.8 OD, was induced with 0.2 mM isopropyl β-D-1-thiogalactopyranoside (IPTG) and incubated further at 22 °C for 16 h. The cells were pelleted and suspended in a lysis buffer containing 20 mM Tris–Cl, 0.5 M NaCl, 1% Triton-X 100, 1X protease inhibitor cocktail, pH 8.0. Cells were disrupted by sonication on ice for 20 min (repeated cycles of 5-sec pulse and 15-sec rest). The crude supernatant was collected from the lysate by centrifugation at 12000g for 10 min, further clarified using 0.2 µm filter and loaded on 5 ml Ni-NTA Sepharose high-affinity pre-packed column (His Trap^TM^, GE healthcare, Sweden), which was pre-equilibrated with binding buffer (20 mM Tris-Cl, 0.5 M NaCl, 20 mM imidazole, pH 8.0). The elution of the bound protein was carried out with a linear gradient of imidazole (0-0.5 M) in the elution buffer (20 mM Tris–Cl, 0.5 M NaCl, 0.5 M imidazole, pH 8.0), the eluted peaks were pooled and analyzed on SDS-PAGE. PsC23 protein-containing fraction was collected and imidazole was removed by 10K centrifugal filter (Merck Millipore, Germany) using buffer (20 mM Tris-Cl, 0.1 M NaCl pH 8.0). The subsequent steps of purification and clean-up were done by size exclusion chromatography (Superdex-S75, GE healthcare, Sweden) in a column pre-equilibrated with phosphate-citrate buffer (50 mM, 0.1 M NaCl, 10% glycerol, pH 8.0). Protein concentration was measured by Bicinchoninic acid assay (Pierce^TM^ BCA protein assay kit). The purified PsC23 protein was stored in 50 mM phosphate-citrate buffer at −20°C for further use.

### Antigen binding and cross-reactivity of PsC23

The antigen-binding activities were measured by modified whole-bacterial cell ELISA (22). The binding affinity of PsC23 to ESKAPE pathogens (*P. aeruginosa, K. pneumoniae, S. aureus. E. faecalis,* and *A. baumannii,* with the exclusion of *Enterobacter spp*) was evaluated. The pure culture of each pathogen (1 x 10^6^ CFU) was coated onto ELISA plates in 50 mM bicarbonate coating buffer, pH 9.2 and air-dried. Wells were blocked with 5% skimmed milk (Sigma-Aldrich, USA) in PBST overnight at 4°C. 100 µL of the purified PsC23 (10 µg/mL in PBST) was then added to each well and incubated further (1h at 37°C). Wells were washed thrice with PBST and monoclonal primary antibody (anti-his-tagged mouse antibody, 1:5000 diluted, Sigma-Aldrich, USA) was added and incubation was continued for 1 h at RT. Secondary polyclonal anti-mouse-HRP-conjugated antibody (Sigma-Aldrich, USA, 1:2500 diluted) was then added to each well and kept for 1h at RT. PBS without primary antibody was used as a control. The plate was washed thrice with PBST and twice with PBS, and 0.05 mM TMB substrate was added. The reaction was terminated by 1% sulfuric acid when the desired intensity of the blue color was developed, which was then measured at 450 nm. Since PsC23 is an antibody fragment, it should have specificity towards a particular pathogen. To check the neutralizing specificity, a purified PsC23 antibody (25 µg/mL) was incubated with *Pseudomonas aeruginosa* as well as different bacteria (*Enterococcus faecalis, Staphylococcus aureus, Klebsiella pneumoniae, Acinetobacter baumannii, Salmonella enterica,* and *Staphylococcus haemolyticus*) at 1 x 10^6^ CFU/mL in MH broth at 37^0^C and growth (OD 625 nm) was measured after 8 h.

### Minimal Inhibitory Concentration (MIC) and pathogen-specificity and time kill kinetics of PsC23

Minimal Inhibitory Concentration (MIC) can be defined as the lowest concentration of antimicrobial agent required for the inhibition of the bacterial growth (23). We determined the MIC-50, 90 and 99 of PsC23 for both *P. aeruginosa* strains (ATCC 27853 and MCC 50428). Antimicrobial activity was determined following the instructions recommended by Clinical and Laboratory Standards Institute (CLSI) guidelines (24). Briefly, the purified PsC23 was diluted (50 µg/mL-1.625 µg/mL) in 200 µL MH broth. Overnight grown culture of each strain at 0.5 McFarland standard suspensions was prepared in saline and 1 x 10^6^ CFU/mL bacteria was added to each well. PBS without PsC23 antibody, was taken as a negative control. The plate was incubated for different time periods at 37°C, 400 rpm, OD 625 nm was recorded and colonies (24 hrs) were counted after plating a suitable dilution on MH agar. Time-kill assays of the purified PsC23 for both *P. aeruginosa* strains were carried out with suitable modifications as described earlier (25). Various concentrations 1X (25 µg/mL), 2X (50 µg/mL), 4X (100 µg/mL) times MIC-99 of PsC23 were inoculated with 1 x 10^6^ CFU/mL of both *P. aeruginosa* strains and evaluated for their inhibitory activity. At time intervals 0, 2, 4, 8, and 24 h, 10 µL samples were withdrawn from each well and plated on MH agar to determine the CFU/ml.

### Adaptive resistance studies on *P. aeruginosa* 27853 and effect of the efflux channel blocker on clinical resistant *P. aeruginosa* MCC 50428

ATCC reference strain *P. aeruginosa* 27853 was grown in presence of a panel of drugs (Piperacillin, Meropenem, Gentamycin, Cefepime, Ciprofloxacin) aerobically and anaerobically (by overlaying mineral oil) and OD600 was recorded after 24 hrs.

*P*. *aeruginosa* 27853 and *P. aeruginosa* MCC 50428 was grown in MH broth at 37^0^C in presence of 1. PsC23 (25 µg/mL) + carbapenem (8 µg/mL) or 2. carbapenem (8 µg/ml) + Phe-Arg-β-naphthylamidedihydrochloride (PAβN) (20 µg/mL) which is a broad-spectrum efflux inhibitor commonly used for drug efflux characterization (26) 3.Meropenem 4.Meropenem+ PAβN 5.Meropenem+ PAβN+PsC23 antibody and OD620 was monitored at various time intervals for 24 hr.

### Effectiveness of combination therapy of PsC23 and meropenem *P. aeruginosa* MCC 50428 by checkerboard analysis

To evaluate the effective combination of meropenem and PsC23 antibody in reversing resistance in meropenem resistant *P aeruginosa* MCC 50428, 10^6^ cells were grown under various combinations of meropenem (32µg/ml - 0.5µg/ml) and PsC23 antibody (50µg/ml - 0.79µg/ml) at 37^0^C for 24 hrs and optical density was recorded periodically. A checker board was made to indicate the probable combinations of these two drugs that were most effective in inhibiting the growth.

### PsC23 target identification by mass spectrometry and functional characterization

The *P. aeruginosa* cells were lysed by sonication on ice in a lysis buffer (protease inhibitor cocktail, SDS (2%), β-mercaptoethanol (50 mM) and supernatant was resolved on 15% SDS-PAGE and proteins were electro-transferred to nitrocellulose membrane for Western Blotting. The membrane was blocked with 5% skimmed milk (1hr), target was then probed by Histidine tagged PsC23 VHH (10 µg) followed by washing and addition of a horseradish peroxidase (HRP)-conjugated anti-his-IgG secondary antibody. After addition of the substrate 3’ Diaminobenzidine (DAB), the immuno-detected band was found to be around 33 kDa (Fig. 5). For confirmation of the target, the corresponding band in SDS-PAGE was excised and processed for in gel trypsin digestion. The peptides were first separated by Liquid chromatography on ACQUITY UPLC system using C18 column. Resulting peptide fractions were directed to electrospray ionization MS/MS (ESI-MS/MS) on Q TOF (Agilent) mass spectrometer and raw data was acquired and processed by an opensource software Morpheus. The protein sequence database of *Pseudomonas aeruginosa* (FASTA) was downloaded from Swiss-prot and used for identification of the peptides and proteins. ATCC reference strain 27853 was grown in a minimal medium containing 40mM each of C4 carbon sources succinate or fumarate in anaerobic conditions to activate the glyoxylate shunt that is fed by C4 carbon sources (27) at 37^0^C with and without PsC23 antibody and growth was monitored for 48 h for functional characterization of the target.

### Role of PsC23 in inhibition of drug efflux by nile red assay

We investigated the role of PAβN and PsC23 in the efflux mediated resistance by a fluorescent dye, nile red. Anaerobically grown overnight grown *P*. *aeruginosa* MCC 50428 (OD600 1.0) was incubated with nile red (5µM) and anti PsC23 antibody (25µg/ml) or PAβN (20µg/ml) separately for 4 hours. Cells were washed and fluorescence intensity was measured (excitation:552, emission:636 nm) following which 100µl sterile succinate and fumarate (100mM) was added in each tube, to activate the glyoxylate shunt and the efflux mechanism, followed by incubation at RT for 15-20 min. Cells were pelleted, washed and fluorescence intensity was measured to quantitate the nile red remaining in the cells.

### *In vivo* efficacy of PsC23 and meropenem against *P. aeruginosa* MCC 50428

These experiments were carried in collaboration with PRADO Preclinical and Research Facility, according to the guidelines and approval of the Institutional Animal Ethics Committee Pvt. Ltd Pune, India. Pathogen free 6-8 weeks old BALB/c mice (female) were housed in polypropylene cages and maintained under controlled conditions (23°C –25°C, 50–55% humidity and 12 h dark and light cycles). They were fed with chow diet and water *ad libitum*. Before infection, neutropenia was induced in these mice by injecting two intra-peritoneal doses of cyclophosphamide at day - 4 (150 mg/kg) and day −1 (100 mg/kg). On the day of infection, an overnight grown culture of meropenem resistant *P. aeruginosa* MCC 50428, 0.5 McFarland standard suspension (equivalent to a density of 10^8^ CFU/mL) was prepared in a sterile lactate ringer’s solution. Infection was induced by injecting 50 µL bacterial suspension into tail vein of mice using 27-guage-needle (28).

A total of five groups (*n*=4) of mice were used. One hour post infection mice received various treatments for 72 h as described in Supplementary table 1 and survival of the control and treated animals was monitored. Heparinized blood samples were collected aseptically after 24 h, centrifuged at 2,000 x g for 5 min and a suitable dilution was plated on MH agar, and incubated at 37^0^C, 24 h for determination of the bacterial load (CFU/mL). The 5^th^ group of mice was administered purified PsC23 (equivalent to 5 mg/kg of body weight) without inducing any infection and examined for the preliminary signs of toxicity. All experiments were approved by the ethics committee with all relevant ethical regulations for animal testing.

### Statistical analysis

The statistical analysis of data was done using GraphPad prism (V5). Student t-test, Analysis of variance (ANOVA) and Bonferroni posttest were applied where ever it was appropriate. Statistical significance is indicated at *P* < 0.005.

## Results

### Construction of *P. aeruginosa* VHH library and isolation of the neutralizing hits

An Indian dromedary camel was immunized with inactivated clinical *P. aeruginosa* MCC 50428 strain and boosted twice every two weeks for one and half months. When a high titer was obtained after second booster dose, 60 mL heparinized blood was collected from camel. PBMCs were isolated and VHH fragments encoding open reading frame were amplified using specific primers (supplementary methods 1) and ligated to the pADL-23c phagemid vector. Freshly ligated VHH-vector constructs were transformed into *E. coli* ER 2738 by electroporation, a library (CIL-Ps) of 2×10^9^ clones was obtained. Colony-PCR screening revealed that over 90% of these colonies contained VHH gene constructs (Fig.1a). CIL-Ps phage display library was enriched against the whole-cell *P. aeruginosa* using 3 rounds of bio-panning. Randomly selected individual binders were inoculated separately in a 96 deep well plate and screened for binding by phage-ELISA (Fig. 1b). Colonies that gave 3-5 folds more reading than negative control were identified as optimal binding hits. 3-8 such hits were isolated from each plate and categorized into strong, moderate, and non-binders. The strongest binders were tested for their neutralizing activity by leaky secretion method developed in house. Out of the 32 binders, five clones namely, PsC11, PsC23, PsC30, PsC56 and PsC79 were found to have a neutralizing activity (Fig. 1c), all except PsC56 inhibited the growth of *P. aeruginosa*. The uniqueness of the neutralizing hits was determined by RFLP. Three of the neutralizing hits PsC11, PsC23 and PsC56 showed a unique restriction digestion pattern, while two clones namely PsC30 and PsC79 were found to be identical (Fig. 1d). One of the clones PsC23 that distinctly inhibited both strains of *P. aeruginosa* was identified as a hit for further characterization.

### Cloning, expression and purification of PsC23

PsC23 VHH gene was sub-cloned into pET28c (+) expression vector, transformed into *E. coli* C41(DE3) and induced with IPTG (0.2M) for heterologous expression. This molecule was 6X Histidine-tagged at N-terminal to enable the ease of purification and characterization. The soluble expression conditions were optimized and highest production was observed in shake flasks at 180 rpm, 22°C for 16 h. Cells were lysed by sonication, the crude supernatant lysate was passed through immobilized metal ion affinity chromatography (IMAC) using His Trap^TM^ pre-packed column. Elution was carried with a linear gradient of imidazole, PsC23 protein was eluted as single peak (Peak 3) at 0.25 M imidazole (Fig. 2a). Eluted fractions were pooled and concentrated after removal of imidazole using 10K centrifugal filter and the yield was 3-4 mg/L. The final purification and clean-up were done by gel filtration chromatography using Superdex-75 resin. A single band of 15 kDa obtained on SDS-PAGE (Fig. 2b) was ascertained to be of more than 95% purity by visual inspection of the gel.

**Fig. 2.**
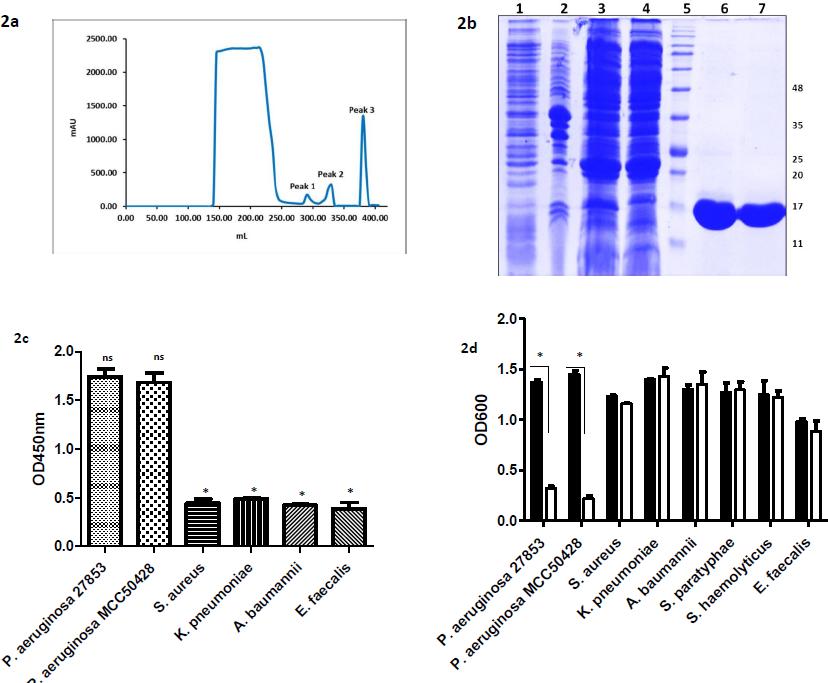
**A.** Ni-Sepharose (His Trap^TM^) affinity chromatography purification of PsC23: Crude lysate supernatant was loaded on a 5 ml pre-packed column and eluted with a linear-gradient range of 0 - 0.5 M imidazole; PsC23 eluted out at peak 3 corresponding to 0.25 M imidazole. **B**. SDS-PAGE analysis of eluates from the chromatographic purification steps of PsC23: **Lane 1-**Uninduced lysate, **Lane 2-** Induced lysate supernatant**, Lane 3-** Peak 1 eluate from Ni-Sepharose **Lane 4-** Peak 2 elute from Ni-Sepharose, **Lane 5-** Molecular weight marker, **Lane 6-** Peak 3 eluted (PsC23), **Lane 7-**Superdex S-75 eluted (PsC23). The molecular weights are indicated on the right. **C.** Quantitation of binding specificity and cross-reactivity of purified PsC23 by whole-cell ELISA to both strains of *P. aeruginosa* and other pathogens from the ESKAPE group. **D.** Specificity of the neutralization action of PsC23 as quantitated by their effect on the growth of different human pathogens.

### PsC23 binds specifically to *P. aeruginosa* and is not cross reactive to other pathogens of ESKAPE group

The binding ability and cross-activity of purified PsC23 to the ESKAPE group pathogens was evaluated using whole-bacterial cell ELISA. The intensity of binding in terms of OD 450 was higher for both strains, ATCC 27853 and MCC 50428 strains of *P. aeruginosa* as compared to other bacteria. This indicated that PsC23 has a high binding specificity to *P. aeruginosa* and has a nominal cross-reactivity with other bacteria (Fig. 2c). As PsC23 demonstrated a neutralizing activity against *P. aeruginosa*, we also examined this effect on other bacteria associated with human health. Purified PsC23 (25µg/ml) inhibited *Pseudomonas* sp. within 8 h but did not significantly affect the growth of other tested pathogens (*Enterococcus faecalis, Staphylococcus aureus, Klebsiella pneumoniae, Acinetobacter baumannii, Salmonella enterica, Staphylococcus haemolyticus*) (Fig. 2d). This indicated that the PsC23 was very specific and did not have much off target activity. Thus, it could be used for development of *P. aeruginosa* specific targeted therapeutics and diagnostics and it would not disrupt the patient microbiota when administered internally.

### PsC23 was effective at a dose of 25 µg/mL against two different strains of *P. aeruginosa*

The MIC of PsC23 for two strains of *P. aeruginosa,* ATCC 27853 and clinical resistant strain MCC 50428 was evaluated *in vitro* and growth monitored periodically. MIC-50, 90 and 99 that correspond to growth inhibition by 50%, 90% and 99% were found to be 6.25, 12.5, and 25 µg/mL respectively for both these test strains. This suggests that the purified antibody fragment (PsC23) is efficacious irrespective of the resistance profile of the strain and might be effective against other clinically resistant *P. aeruginosa* as well.

### A combination of PsC23 and carbapenems is more effective than PsC23 alone for control of drug resistant *P. aeruginosa*

We performed time-kill assay of PsC23 against both strains (ATCC 27853, MCC 50428) of *P. aeruginosa* at 1X (25 µg/mL), 2X (50 µg/mL) and 4X (100 µg/mL) of the effective dose and monitored the number of viable cells over the period 2, 4, 8, and 24 h (Fig. 3a, 3b). The difference in the growth profiles was not significant at these concentrations indicating that a threshold might have been obtained at 25 µg/mL, the most pronounced effect was seen within 2 h after which the cells recovered and a gently upward sloping curve was observed. Moreover, the effect of the antibody seems to be maximum around 2 h, after which the population seems to recover slightly. Surprisingly, when the clinical *P. aeruginosa* MCC 50428 strain was treated with PsC23 (25 µg/mL) and carbapenems (8 µg/mL meropenem and imipenem), a bactericidal effect was observed to be more pronounced and no further growth was seen after 4 h (Fig. 3c, 3d). We investigated this phenomenon further by using different combinations of PsC23 and meropenem by a checker board analysis on the *P aeruginosa* MCC 50428 strain (Fig 4). Meropenem resistance was completely reversed when only 4 µg/ml of meropenem was used with 50 µg/ml of the PsC23. When the amount of meropenem was increased to 8, 16 and 32 µg/ml, the PsC23 dose went down to 12.5, 6.25 and 1.58 µg/ml. These data give an idea of the range of concentrations the PsC23 VHH can be used to reverse meropenem resistance, the optimal ratio of which can be fixed depending on the druggable properties of the combination and the economy factor.

**Fig. 3.**
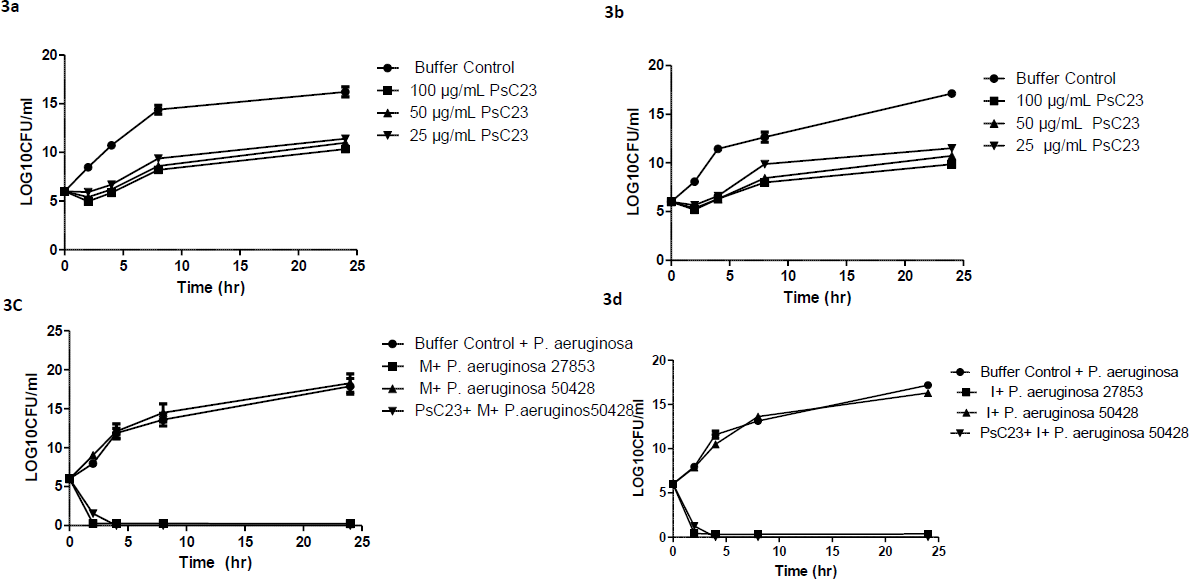
Growth kinetics of *P. aeruginosa* after treatment with PsC23 over a period of 24 h. **A .**Effect of 25 µg/mL, 50 µg/mL and 100 µg/mL (1X MIC, 2X MIC, and 4X MIC-99, respectively) PsC23 on the growth of a reference *P. aeruginosa* ATCC 27853 strain compared to a control with no treatment **B.** Effect of 25 µg/mL, 50 µg/mL and 100 µg/mL (1X MIC, 2X MIC, and 4X MIC-99, respectively) PsC23 on the growth of a clinical *P. aeruginosa* MCC 50428 strain resistant to meropenem (M) compared to a control with no treatment**. C.** Growth kinetics of *P. aeruginosa* ATCC 27853 and clinical MCC 50428 in the presence of meropenem (8 µg/mL), PsC23 (25 µg/mL), and a combination of PsC23 with meropenem. **D.** Growth kinetics of *P. aeruginosa* ATCC 27853 and clinical MCC 50428 in the presence of imipenem I (8 µg/mL), PsC23 (25 µg/mL) and a combination of PsC23 with imipenem. ANOVA followed by Bonferroni post-test indicates a decline in growth of carbapenem-resistant *P. aeruginosa* MCC 50428 in presence of antibody plus meropenem or antibody plus imipenem compared to meropenem or imipenem alone (\**P*<0.005).

**Fig.4.**
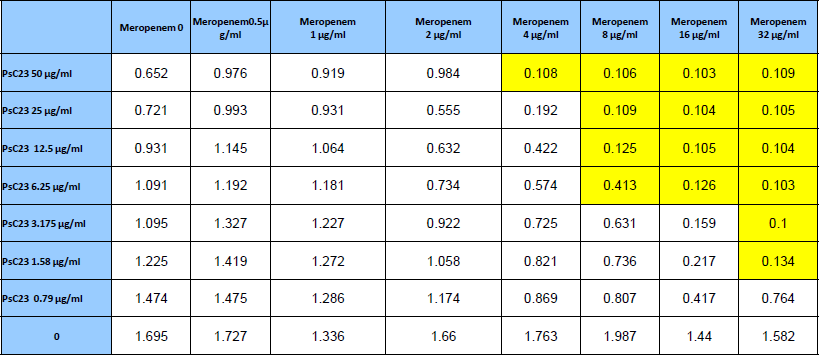
Checker board analysis to evaluate the combination of meropenem and PsC23 on the growth of *P.aeruginosa* MCC 50428.

### The target of PsC23 is a component of the C4 dicarboxylate transporter as identified by mass spectrometry

Mass spectrometry analysis identified a component of C4 dicarboxylate transporter as a binding target of PsC23 VHH with high confidence (Fig.5). This transporter is usually present on the cell membrane of *P. aeruginosa* and other bacteria and facilitate the uptake of C4 carbon sources that is fed into the glyoxylate shunt that is operative under conditions of oxidative stress in the host. Under oxygen limiting conditions when the TCA cycle is not fully operative, glyoxylate shunt is used for energy generation and anabolic activities (27) and has a critical survival value in pathogenic bacteria. It is present in bacteria but is absent in the eukaryotic host, making it a suitable target for drug development.

**Fig.5.**
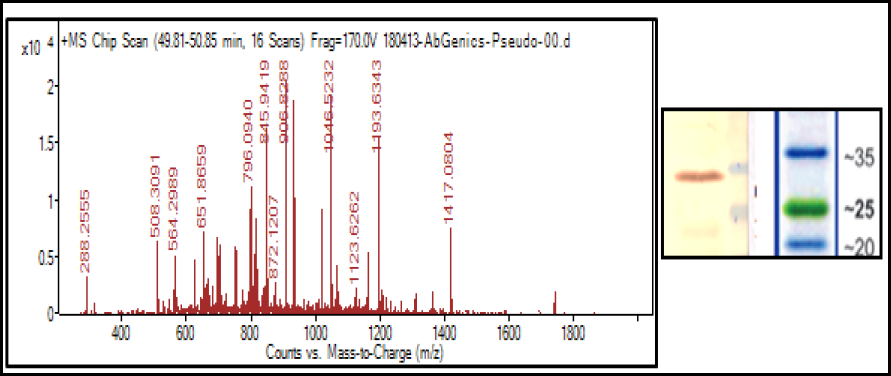
**A.** Representative total ion current image of the mass spectrometry-based analysis of antibody target **B.** Western blot showing localization of PsC23 target protein band.

### Active efflux is responsible for resistance and PsC23 inhibits growth of *P. aeruginosa* in with C4 as sole carbon source

The accumulation of the fluorescent dye nile red in *P.aeruginosa* MCC 50428 when grown in the presence of succinate as the sole carbon source was used to ascertain if the resistant strain accumulated less dye and if the effect could be reversed by blocking the C4 dicarboxylate transporter by PsC23 or efflux by PAβN. When succinate was added to the anaerobically grown culture, the dye accumulation was observed to be 50% less that the control. This effect was reversed considerably by addition of both the PsC23 as well as PAβN (Fig 6a) indicating that both the compounds interfere with the efflux of the dye. The blocking of C4 dicarboxylate transporter by PsC23 should be optimally assessed if *P. aeruginosa* is grown in minimal media supplemented with C4 carbon sources anaerobically and the growth monitored periodically. When *Pseudomonas aeruginosa* was grown in a minimal medium supplemented with C4 carbon sources succinate (Fig. 6b) or fumarate (Fig. 6c), normal growth was seen. When PsC23 (25µg/ml) was added the growth was found to be inhibited in both conditions, which could be probably due to blockage of the import of C4 carbon substrates by PsC23 that leads to the blockage of efflux by a yet uncharacterized mechanism.

**Fig 6.**
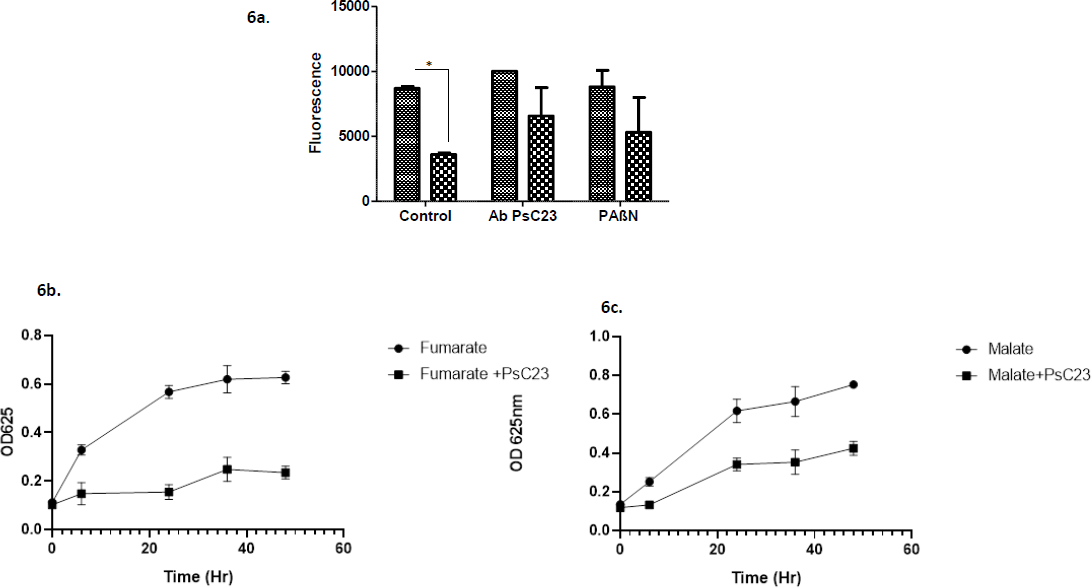
Growth of *P.aeruginosa* MCC50428 in minimal media supplemented with C4 carbon sources and effect of PsC23 and PAβN on drug efflux and growth profiles **a**. Effect of PAβN, and PsC23 on the accumulation of nile red dye in in the presence of succinate as the sole carbon source **b,c**. Effect of PsC23 on the growth of *P.aeruginosa* MCC50428 under anaerobic condition in minimal medium supplemented with b. Succinate only (•) and in combination with PsC23(▪). c. Fumarate only (•) and in combination with PsC23 (▪).

### PsC23 reverses efflux mediated resistance in a carbapenem resistant *P. aeruginosa*

To further evaluate the role of C4 dicarboxylate transporter in resistance, we studied the effect of PsC23 on *P. aeruginosa* ATCC 27853 grown under anaerobic condition where the glyoxylate shunt is activated. When ATCC 27853 strain was grown in presence of 5 different drugs, aerobically as well as anaerobically, a reduction in the efficacy of all the drugs was seen under anaerobic condition, indicating that it starts acquiring adaptive resistance to them (Fig 7a). This was a surprising finding and we wanted to investigate the mechanism of such widespread resistance and if the same was operative in the clinically resistant strain MCC 50428. As one of the mechanisms of acquired resistance to multiple drugs is through active efflux of drugs, we studied the effect of PsC23 on drug efflux in *P. aeruginosa* MCC 50428 by comparing the change in resistance profile of carbapenems with efflux blockers. *P. aeruginosa* was incubated with either a. carbapenems, meropenem and imipenem (8 µg/mL) b. carbapenems (8 µg/mL) and PsC23 (25 µg/mL), c. carbapenems (8 µg/mL) and PAβN (20 µg/mL), and d. PAβN (20 µg/mL) respectively and growth was monitored for 24 h. Carbapenems had no effect on the resistant strain as seen in earlier experiments whereas addition of PAβN reversed this effect within 2 h. The similar inhibition pattern for resistant strain was also observed when meropenem or imipenem and PsC23 were used in combination (Fig.7b, 7c). It indicates a possible efflux-mediated resistance to carbapenem that is reversed by PAβN or PsC23. There is a similarity in the mode of action of PAβN and PsC23, which is the blocking of drug efflux that prevented carbapenem efflux from *P. aeruginosa*. The other possibility may be that antibody fragment targets an energy transduction mechanism that energizes an efflux pump by a yet uncharacterized mechanism (Fig 8), otherwise PsC23 would not have inhibited the growth of resistant *P. aeruginosa* when applied with carbapenems.

**Fig.7.**
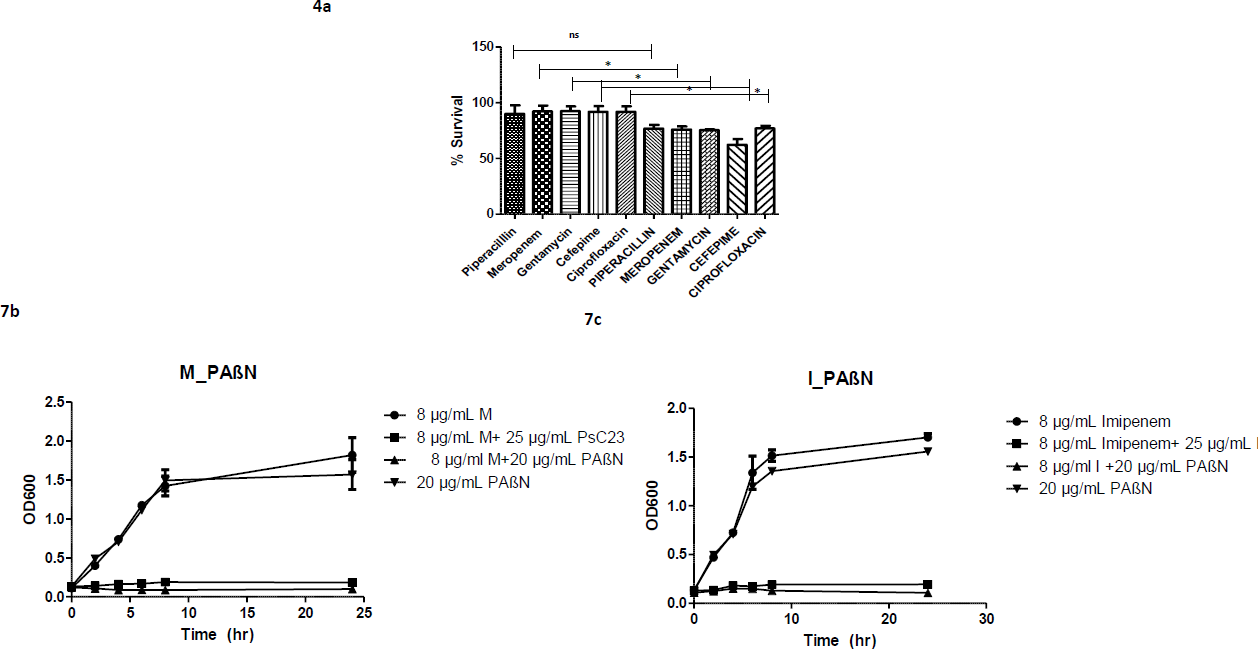
**a.** Effect of anaerobic conditions on the drug resistance profile of *P.aeruginosa* ATCC 27853 on five different classes of drugs. The percentage survival in aerobic conditions is indicated on the left in small case and those under anaerobic conditions in uppercase on the right. **Fig.7b.** Effect of meropenem M (8 µg/mL), PAβN (20 µg/mL), and PsC23 (25 µg/mL) individually and in combinations (meropenem with PsC23 or meropenem with PAβN) on the growth of meropenem resistant *P. aeruginosa* MCC 50428. **Fig.7c.** Effect of imipenem I (8 µg/mL), PAβN (20 µg/mL), and PsC23 (25 µg/mL) individually and in combination (imipenem with PsC23 and imipenem with PAβN) on the growth of carbapenem-resistant *P. aeruginosa* MCC 50428. ANOVA followed by Bonferroni posttest indicates a decline in growth of carbapenem-resistant *P. aeruginosa* MCC 50428 in presence of antibody plus meropenem or antibody plus imipenem or antibiotics plus PAβN compared to meropenem or imipenem or PAβN alone (\**P*<0.005).

**Fig.8.**
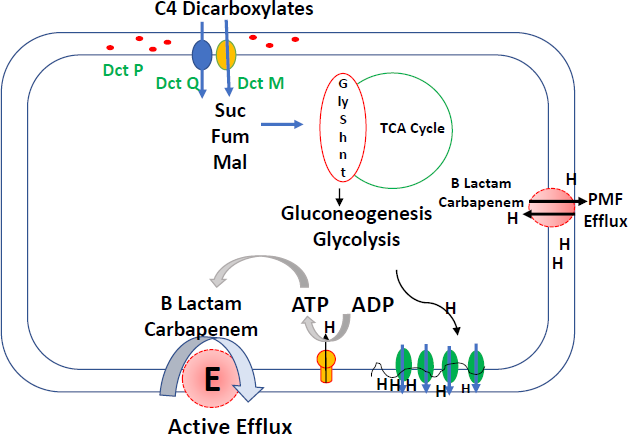
Hypothetical mechanism of action of PsC23 and the role of glyoxylate shunt in efflux mediated drug resistance. The PsC23 might be binding to either the Dct P, Q or M subunits of the C4 dicarboxylate transporter, thereby blocking the uptake of C4 metabolites that are fed into the glyoxylate shunt that energizes active efflux or is involved in a proton motive force antiport based efflux that might result in adaptive resistance to multiple drugs.

### PsC23 and meropenem clears a carbapenem resistant *P aeruginosa* MCC50428 systemic infection in a neutropenic BALB/c mouse

Before advancing the PsC23 to *in vivo* studies its druggable properties were evaluated. It was found to be stable in human serum for 24 hrs (suppl fig 1a) and did not have a toxic effect on HEK293 cells even at 100µg/ml as evaluated by MTT assay (suppl. fig 1b) or analysis of hemolytic activity (suppl. fig 1c) To evaluate the *in vivo* efficacy of PsC23, its effect was studied in BALB/c mice with systemic *P. aeruginosa* infection. One hour post induction of infection with *P. aeruginosa* MCC 50428, various treatments of PsC23, meropenem and their combinations were administered intravenously as mentioned in the Suppl. Table 1. Heparinized blood was withdrawn from the tail vein after 24 h and the pathogen load was counted by plating. A significant increase in bacterial counts (approx. 12-logs) in control mice was noted, suggesting a rampant systemic infection. Similarly high bacterial counts were seen in mice administered with 5 mg/kg body weight dose of meropenem, (suppl fig. 2a). Both these groups of mice appeared sick and morbid; their movements were sluggish. Treatment of PsC23 alone and in combination with meropenem reduced the colony counts by 8-logs in the former and 10-logs in the later compared to control mice. The lower bacterial counts in later might be due to synergistic effect of both meropenem and PsC23. The untreated mice showed 100% mortality within 72 h due to a high bacterial load in blood (suppl fig. 2b). On the other hand, treated mice showed no mortality till 72 h. The 5^th^ group of mice that were administered with PsC23 equivalent to 5 mg/kg body weight without infection also survived for 72 h (suppl fig. 2b) indicating non-toxic property of the VHH. PsC23 is a new molecule with a unique target and in combination with meropenem might be useful for controlling carbapenem resistance in *P aeruginosa* if the two are co administered together.

## Discussion

Control over drug resistant bacteria can be achieved by regulating the use of antibiotics in addition to constantly developing newer molecules having unique modes of action. Among the alternative approaches being developed for treatment of MDR pathogens, antibodies which can inactivate these pathogens, their virulence factors or toxins are expected to have a major clinical impact in coming years. With its demonstrated efficacy for treatment in various types of cancers and autoimmune diseases along with a robust scientific and regulatory framework, antibodies promise to be the most impactful technological advances in the anti-infective space (29). In order to develop antibody based therapeutic for treatment of drug resistant *P. aeruginosa*, we used novel form of antibodies derived from camels (VHH) which are directed towards the surface targets of this pathogen and have demonstrated their use as an adjunct therapy when combined with carbapenems to control the resistant *P. aeruginosa*. We used a novel strategy of leaky secretion analysis for isolating neutralizing VHH antibody fragments against the surface antigens of *P. aeruginosa* without any prior target information. One of the hits (PsC23) isolated from an immunized camelid antibody library having a good neutralizing activity was cloned and expressed in *E coli*, the final yield of a purified VHH protein obtained in our study was about 3.5 mg/L which was higher than previously reported for soluble VHH fragments (30). Earlier studies also used neutralization based approaches using purified VHH antibodies against food-borne pathogen *Listeria monocytogenes* by targeting Internalin B surface protein (31) as well as human viruses including influenza and hepatitis C virus (HCV), respiratory syncytial virus (RSV), human immunodeficiency virus HIV-1, and some enteric viruses (32). The uniqueness of our approach lies in the fact that neutralization assay was done with the unpurified VHH released in the supernatant that allowed for high throughput analysis of multiple clones rapidly. Antibody based biologicals have a key property of specificity that sets it apart from small molecules that can be used to develop target specific therapy as well as accurate diagnostic tests. There are few reports on *in-vitro* cross-reactivity of VHH fragments against viruses (33, 34) and bacteria (35). However, the specificity of the PsC23 binding to *P aeruginosa* and reproducibility of this technique indicates the robustness of our screening strategy, which can be used to quickly isolate antibodies against other pathogens as well.

Furthermore, this antibody fragment was found to be highly effective for both strains of *P. aeruginosa*, one of them being clinically resistant to multiple drugs. The target was revealed to be a component of C4 dicarboxylate transporter by mass spectrometry that is responsible for the uptake of C4 metabolites under oxidative stress that is fed into the glyoxylate shunt. Role of this target in efflux mediated resistance not been reported earlier. As glyoxylate shunt is present in bacterial pathogens and not in the human host, it opens the possibility of developing newer drugs targeting this pathway. However, the growth kinetics revealed that complete neutralization was not achieved which could be due to the presence of persisters in the population that survived but did not multiply significantly (36). Complete killing of the persisters was achieved only when the combination of carbapenem and PsC23 was used. Similar kind of synergistic effect of meropenem with panobacumab IgM antibody has been reported in neutropenic mice for meropenem resistant *P. aeruginosa* (28). This finding is significant in the light of reports that around 45% of *P. aeruginosa* strains are meropenem resistant in critical care setting in North India (37) and a combination of PsC23 with meropenem can be used to treat these cases.

Our study also suggests a role of drug efflux in resistance as it is reversed by efflux blockers. These adaptive mechanisms are common in *P. aeruginosa* and are used by the pathogens when lodged in the human hosts like in cystic fibrosis patients (38). The *in vitro* restoration of carbapenem efficacy in a resistant clinical strain MCC 50428 upon treatment with PsC23 was a surprising finding with tremendous practical therapeutic implications. Blockage of the efflux pump directly or indirectly may lead to drug accumulation inside the bacterial cell resulting in a bactericidal effect. The reported efflux systems involved in antibiotic resistance in *P. aeruginosa* belongs to the resistance-nodulation-division (RND) family consisting predominantly two types of efflux pump systems the MexAB-OprM, and MexXY-OprM that confer resistance to multiple antibiotics (38). Our results indicate that the glyoxylate shunt might have a role to play in energizing these transporters or some new yet unidentified transporter may be involved. To study the possible mechanism of the reversal of carbapenem resistance *in vitro* when meropenem was applied with PsC23 VHH, we used PAβN, a broad-spectrum efflux blocker and observed growth inhibition in its presence indicating that PsC23 might be affecting the activity of the membrane transporter involved in drug efflux. Purified PsC23 was also found to be non-hemolytic, non-toxic to mammalian cells and stable in human serum at 37^0^C. We then checked the efficacy of the PsC23 alone and in combination with meropenem on the systemic infection murine model infected with carbapenem resistant *P. aeruginosa* MCC 50428. Whereas meropenem had no effect on controlling the *P. aeruginosa* infection, all mice died within 3 days post infection, the PsC23 and meropenem treatment ensured complete survival of mice with 8-10 logs difference in the bacterial counts in blood demonstrating that this molecule is effective *in vivo*. Our results support an earlier observation where significant improvement in the survival of *P. aeruginosa* lung infected mice when treated with combination of various antibiotics with a different antibody Mab166 (39). The blockage of this target by the VHH inhibited the growth of *P.aeruginosa* on C4 substrates, thereby confirming that molecule possibly inhibits the transport of C4 carbon sources. The C4 carbon sources are utilized by bacteria under conditions of oxidative stress that are fed into the glyoxylate shunt. The energy generated by this shunt might have a role in efflux of the drug and blockage of the uptake of C4 metabolites by PsC23 debilitate the pump by a yet uncharacterized mode of action. We studied if efflux could be responsible by studying the trafficking of an indicator dye nile red. Nile red efflux was activated in MCC 50428 after addition of succinate and was reversed by PAβN, as well as by PsC23 (Fig 7a). This supports our hypothesis that PsC23 inhibits the efflux pumps although the mode of action is currently not known. We believe it could be either due to inhibition of NADPH production by disrupting the proton motive force, two mechanisms that need to be explored further. (Fig.8).

Use of the small molecule antibiotics to cure infection disrupts the normal patient microbiome which is followed by immune dysregulation that aggravates the disease severity in critically ill and hospitalized patients (40). One of the advantages of using an antibody for therapy is specificity that is rarely achieved by a conventional small-molecule antibiotic. As this is a narrow spectrum precision antibiotic molecule that will not disrupt patient’s microbiota, it might be particularly helpful for reducing the morbidity in chronically ill patients in hospital settings. Combined with the fact that the target is novel and absent in the mammalian cells, PsC23 is an ideal lead molecule that can be developed as a drug to treat carbapenem resistant *P. aeruginosa*.

## Data availability statement

The data that support the findings of this study are available from the corresponding author Sanjiban K. Banerjee upon request.

## Statements and declaration

All authors have nothing to disclose

## Abbreviations

CDC: Center for Disease Control
ESKAPE: *Enterococcus faecium, Staphylococcus aureus, Klebsiella pneumoniae, Acinetobacter baumannii, Pseudomonas aeruginosa* and *Enterobacter spp*
(MDR): multi drug resistant
MTT: 3-(4, 5-dimethylthiazol-2-yl)-2, 5-diphenyl tetrazolium bromide
MIC: Minimal Inhibitory Concentration
PBMCs: Peripheral blood mononuclear cells
PAβN: Phe-Arg-β-naphthylamidedihydrochloride
sdAb: single-domain antibody
TMB: Tetra-methylbenzidine
VHH: variable heavy chain domain.

## Acknowledgments

This work was partially funded by Bill and Melinda Gates Foundation, Grand Challenges Grant GCE-INDIA/R4/2018/007. We are grateful to Dr. Vijay Satav of Dr. Satav’s pathological laboratories for providing us with the clinical samples of *P aeruginosa* as well as the other pathogens for developing and testing the specificity of the camelid antibodies. Late Mr. Anil Nahar for constant encouragement and funding to build up the infrastructure required for the execution of the project. The authors acknowledge Mrs. Manisha Sabnis, Agastya Suresh, and Rija Nada for the initial development of the phage display and protein purification protocols and Dr. Aditi Ambekar for the critical evaluation of the manuscript.

## Authors contribution

SB formulated the concept and the therapeutic approach and was in charge of the overall planning of the project. AN, PP and MB developed the protein purification strategies and executed the animal studies. MS developed the VHH libraries and identified the hits and conducted the cell-based assays. MK conducted the *in vitro* microbiology assays. All authors read and approved the manuscript.

## Supplementary Figures.

**Fig 1. A.** Serum stability of PsC23 VHH over a period of 24 h at 37°C. **Lane (1-5)** - Time periods of incubations of 0, 2, 4, 8, and 24 h respectively. **Lane 6**-Molecular weight markers, **Lane 7**-Purified PsC23 without serum, **Lane 8**-Human serum without PsC23. Molecular weights are indicated on the right. **B.** MTT assay for evaluation of the cytotoxic effect of purified PsC23 on HEK 293 cell line. The cells were incubated with 1X, 2X and 4X MIC-99 concentration of PsC23 for 24 h and optical density at 570 nm was plotted after the addition of MTT. 1% Triton X-100 which is cytotoxic to the cells was used as a positive control. **C.** Hemolysis assay quantitating the amount of hemoglobin released from RBCs at 1h with 4X MIC-99 concentration of PsC23 on freshly isolated human erythrocytes. 1% Triton X-100 was used as a positive control.

**Fig 2. A.** Bacterial load estimation in blood of BALB/c mice (Log10 CFU/mL) after 24 h of infection with *P. aeruginosa* MCC 50428 without treatment (buffer control), with meropenem (5mg/kg), PsC23 (5mg/kg) and PsC23+ meropenem (5 mg/kg each). **B.** Percentage survival of meropenem resistant *P. aeruginosa* MCC 50428 infected neutropenic BALB/c mice when treated with meropenem (5 mg/kg), PsC23 (5 mg/kg) and PsC23 with meropenem (5mg/kg each) and no treatment (buffer treated) monitored over a period of 72 h.

## Tables

**Supplementary Table 1:**
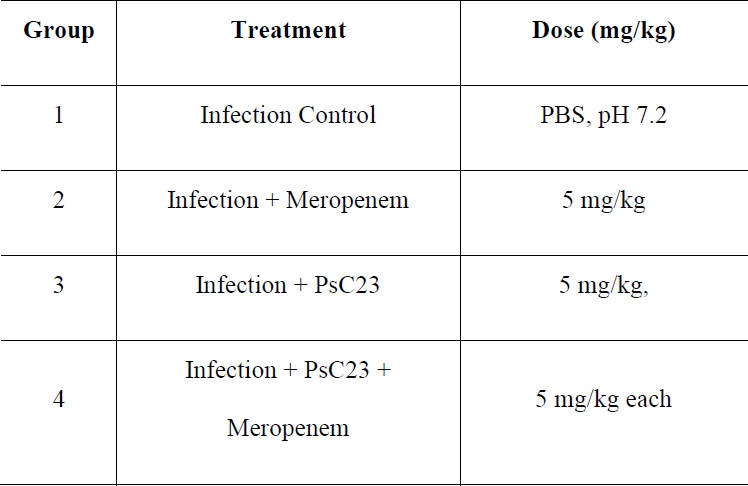
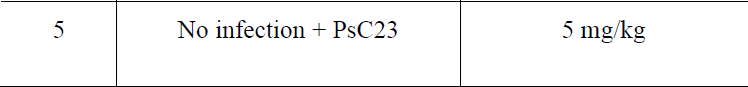
Experimental design to study the effect of PsC23 and meropenem in *P. aeruginosa* MCC 50428 infected neutropenic BALB/c mice.

| Group | Treatment | Dose (mg/kg) |
| --- | --- | --- |
| 1 | Infection Control | PBS, pH 7.2 |
| 2 | Infection + Meropenem | 5 mg/kg |
| 3 | Infection + PsC23 | 5 mg/kg, |
| 4 | Infection + PsC23 +<br>Meropenem | 5 mg/kg each |
| 5 | No infection + PsC23 | 5 mg/kg |

