## Supplementary figures and images for "Reversal of carbapenem resistance in *Pseudomonas aeruginosa* by camelid single domain antibody fragment (VHH) against the C4 dicarboxylate transporter"

### Supplementary Figures

Supplementary fig. 1

1a

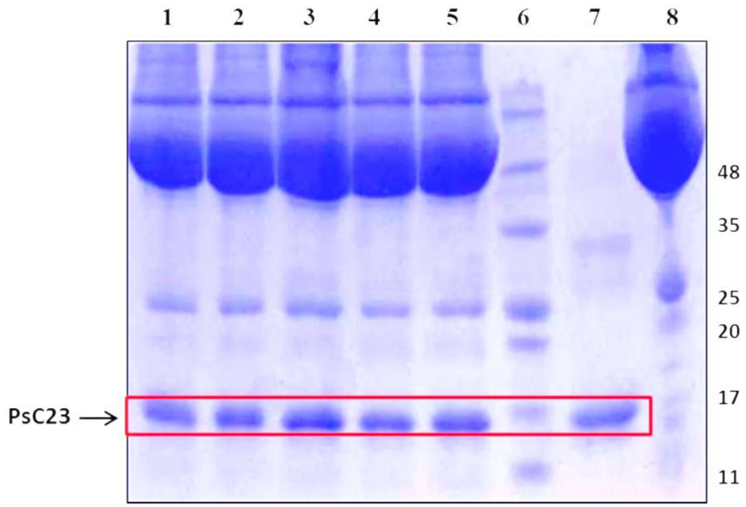


1b

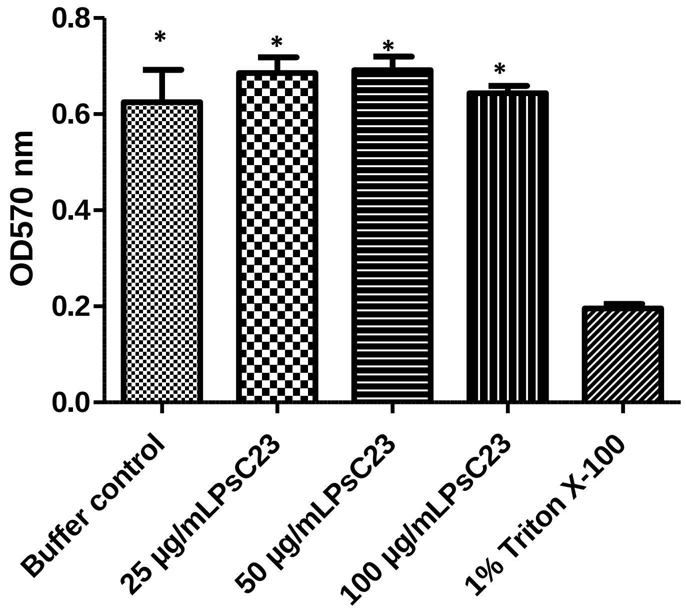


1c

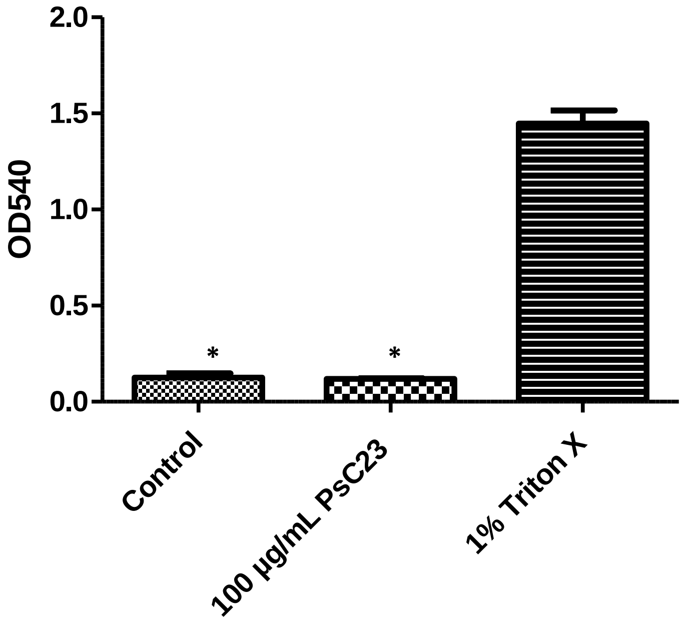

Supplementary fig. 2

2a.

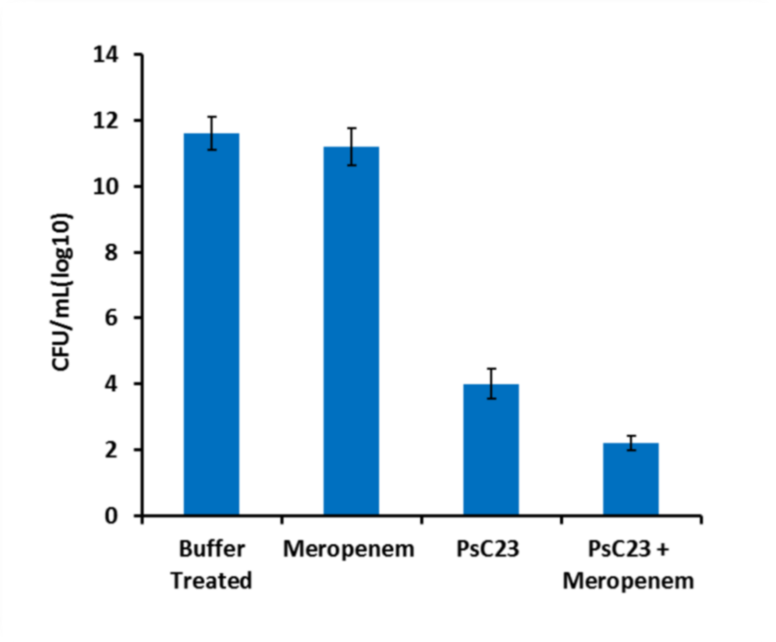


2b.

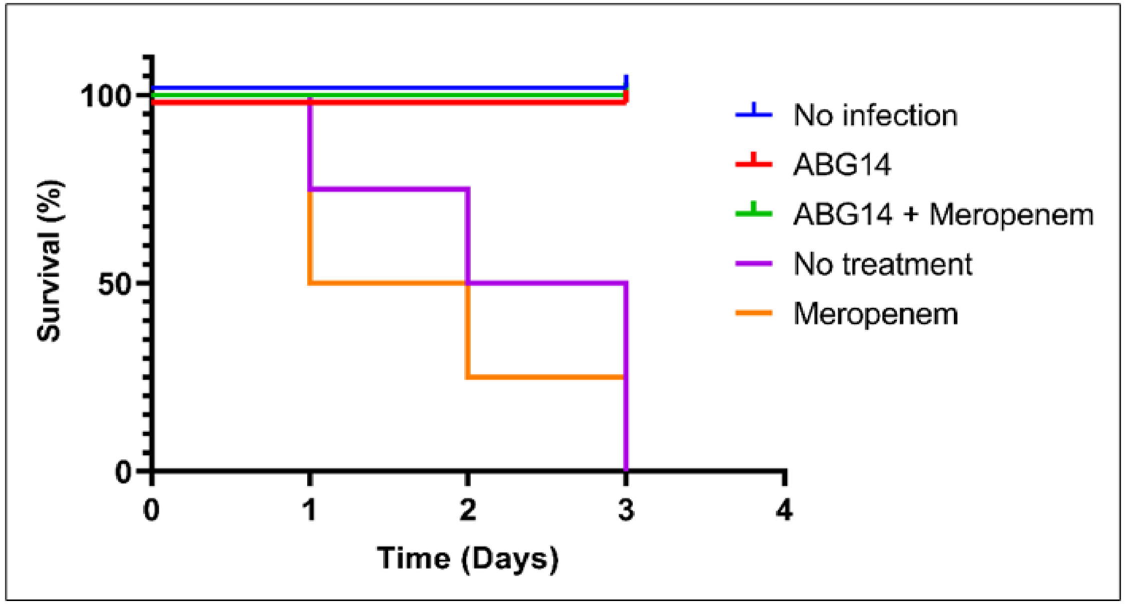
